# Exploring the importance and preference of sugar feeding behaviour of malaria vectors in sugar plantations of southern Malawi

**DOI:** 10.1101/2025.11.06.687027

**Authors:** Kennedy Zembere, Sylvester Coleman, James Chirombo, Rex Mbewe, Julie-Anne Tangena

## Abstract

**Background:** Reliable tools are needed to control opportunistic outdoor biting and resting malaria vectors that remain beyond the reach of indoor targeted interventions. The attractive targeted sugar baits (ATSBs) have demonstrated effectiveness in some settings but have shown limited impact in other areas, likely due to differences in mosquito species’ preferences and the presence of competing natural sugar sources. However, these factors remain poorly understood and understudied. We evaluated sugar-feeding behaviour of Anopheline mosquitoes in a sugarcane plantation area of Chikwawa, Malawi, to generate additional field data that could enhance the efficacy and design of sugar-based vector control tools tailored for malaria endemic regions such as Malawi.

**Methods:** Using three collection tools, CDC Light traps; Prokopack aspirator and the barrier screen, we collected 187 adult anophelines from the Illovo sugar plantations. Collected mosquitoes were subjected to cold anthrone tests in the laboratory to assess the presence of plant sugars in their gut. Additionally, 810 adult *Anopheles gambiae* s.l., reared in the insectary from wild caught larvae, were exposed in an olfactory-driven choice experiment to identify the most attractive available sugar source in the area. Sugar sources included guavas, melon, bananas, mango, marula and sugarcane.

**Results:** Over 40% (n=74) of the collected *Anopheles* mosquitoes-including *An. gambiae* s.l., *An. funestus*, *An. coustani* and *An. tenebrous* were found to have fed on natural sugar sources. For the sugar attractiveness tests for *An. gambiae* s.l., guava was found to be twice as attractive (IRR = 1.97, 95% CI: 1.49-2.62, p < 0.001) as sugarcane (our reference fruit), followed by banana (IRR= 1.68, 95% CI: 1.26-2.24, P < 0.001), then mango, and melon (IRR= 1.49, 95% CI: 1.11-2.01, P= 0.008) and (IRR=1.45, 95% CI: 1.08-1.96, P= 0.014) respectively.

**Conclusion:** Sugar feeding is a key activity for *Anopheles* mosquitoes and presents a potential target for control efforts. In this setting with abundant sugarcane, guava was identified as the most attractive sugar source for *An. gambiae* s.l., followed by banana, mango, and melon, with sugarcane being the least attractive. Understanding local sugar source preferences can help tailor novel intervention strategies to specific environmental contexts.

## Introduction

Insecticide-treated bed nets (ITNs), indoor residual spraying (IRS), and malaria case management using artemisinin-based combination therapies (ACTs) have contributed to worldwide malaria transmission reduction (1–3). Between 2000 and 2015, malaria deaths reduced by over 50% worldwide (4). In the same period, over 663 million cases of malaria were averted with most of this success attributed to ITN use (4). Despite this success, malaria remains one of the leading causes of mortality in Africa, even in areas with good vector control coverage (5). This has, in part, been attributed to the increased level of insecticide resistance and behavioural shifts in vectors and the focus of vector control tools on only indoor biting and resting vectors. Thus, ITNs and IRS use alone cannot achieve complete malaria control and elimination, as opportunistic outdoor biting and resting vectors remain beyond their reach, leading to sustained residual malaria transmission (6,7).

Alternative control tools have been proposed for use in both indoor and outdoor environments to complement existing vector control tools. One promising approach is the attractive targeted sugar bait (ATSB). These traps use a combination of fruits or flower scents (baits), oral toxins and sugar to attract mosquitoes (8,9), leveraging their need for glucose meals (10). Unlike traditional vector control tools that target indoor resting and biting mosquitoes, the ATSB target the entire vector population, including males, and have the potential to reduce the density of both indoor and outdoor mosquitoes (11). Despite promising results from earlier entomological trials in Mali and Israel, recent large-scale trials in Kenya and Zambia (12,13) failed to demonstrate significant epidemiological or entomological impacts of ATSBs, raising critical questions about the context-specific efficacy of ATSBs and the underlying sugar-feeding behaviours of vector populations. Further studies from different ecological settings have thus been recommended (14).

Effective implementation of ATSBs and other tools that exploit sugar feeding behaviour requires thorough understanding of the role of sugar sources in malaria vector ecology, the attractiveness of different sugar sources, and overall mosquito sugar-feeding behaviour. Previous studies have shown that *Anopheles gambiae s.l.* exhibits strong preference for specific sugar sources (15,16). These findings emphasize that local mosquito sugar-feeding behaviour, especially in sugar-rich environments like southern Malawi, can determine ATSB success. While the question of which blood hosts mosquitoes prefer has been extensively studied, in Malawi and across Africa, little is known about the sugar sources most commonly foraged by malaria vectors (16). This study evaluated the anopheline mosquito sugar feeding behaviour as a baseline to understand the importance and preference of sugar sources, providing key insights to guide implementation of vector control tools that exploit sugar feeding behaviour.

## Methodology

### Study area

The study was conducted within the Illovo sugar estate in Nchalo, Chikwawa (− 16.1869; 34.8805), the largest sugarcane producer in Malawi. More details regarding the study area have been discussed elsewhere (17) but in summary, the estate sits within a malaria endemic region in the Shire Valley of Chikwawa district, southern Malawi. *Anopheles arabiensis* and *An. funestus* are the two major vectors in the study area (17). With over 13,000 hectares of sugarcane plantations, the study area relies heavily on extensive irrigation systems during the dry season, a situation that sustains mosquito breeding all year round.

### Testing presence of sugar in the gut of wild caught *Anopheles* mosquitoes

Adult mosquitoes were collected in August 2023 for a period of four weeks, with two consecutive days of collection each week. Mosquitoes were collected using CDC light traps, Prokopack aspirators and barrier screens outdoors next to sugarcane farming plots, focusing on areas near putative breeding sites and surrounding bushes to target newly emerged mosquitoes. It has been shown previously that newly emerged and nulliparous mosquitoes are more likely to feed on sugar sources than older mosquitoes that have previously blood fed or are parous (18). Thus, newly emerged mosquitoes are a key target for sugar feeding exploitation methods. Barrier screens were also set near sugar plantations and near bushes and breeding sites that were close to human habitations to target host seeking mosquitoes that fly towards the households to seek blood or those leaving houses looking for places to oviposit. All mosquitoes were identified to species level. Both male and female *Anopheles* mosquitoes were tested for the presence of sugar.

To determine if the *Anopheles* mosquitoes collected had fed on sugars, we conducted the cold anthrone test which identifies the presence of fructose in the mosquito gut (19). Within 2 hours of collection all *Anopheles* mosquitoes were tested for the presence of fructose before the sugar degraded/digested inside the mosquito gut (20). A 500 ml cold anthrone yellow reagent was prepared by adding 1 gram of anthrone powder to a mixture of 140 ml of distilled water and 360 ml concentrated sulphuric acid in a 1000 ml screw cap bottle. To test for the presence of fructose in mosquito guts, mosquitoes were singly placed in ELISA plate wells (Figure 1). Well A1 was left for positive control with a known sugar (table sugar-sucrose) and B1-H1 were left for negative control with distilled water only.

**Figure 1.**
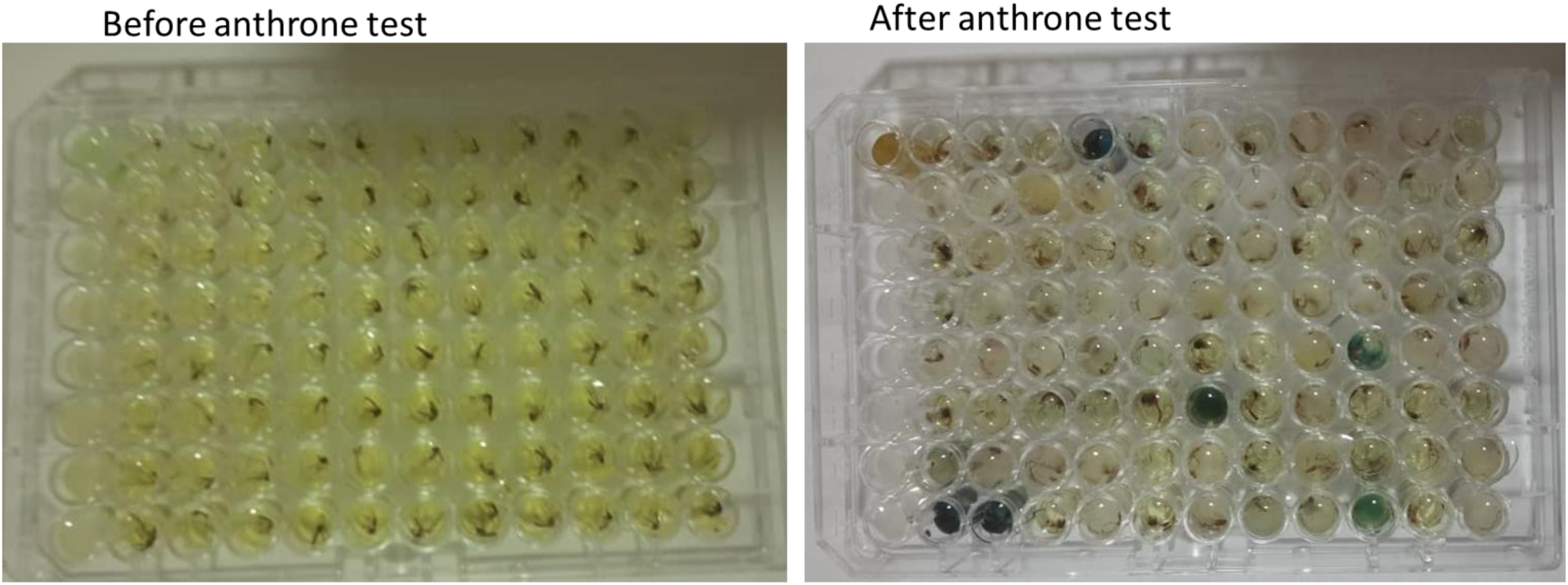
Cold anthrone test. **A**. Mosquitoes in ELISA plates before cold anthrone test. **B**. Mosquitoes that fed on plant sugars changed colour to blue-green after a cold anthrone test.

Whole individual mosquitoes were placed in each well and crushed using separate pestles to release the fructose. A total of 200 ul of cold anthrone reagent was dispensed into each well. The plate was then covered and incubated for 45 minutes at room temperature. After incubation, the results were scored based on the colouration: where yellow (no colour change) was interpreted as negative (no sugar) and blue green colour change indicated a positive result (presence of fructose (Figure 1).

### Sugar choice experiments

Mosquito larvae were collected in the field and reared in the laboratory for the olfactory-driven behavioural assay. Within the Chikwawa sugar plantation area random sampling was done, stratified by proximity to sugar plantation, to select areas for larval surveillance. Stratification was done to maximise larvae presence as it has previously been shown that pollen-rich standing pools of water, associated with the sugar plantations, are potential mosquito oviposition sites and larval habitats for gravid *An. arabiensis* (21). We used probability sampling methods to ensure that all the breeding sites had a chance of being selected. Thus, all breeding sites had equal chances of being selected if they were near sugarcane plantations. We sampled a total of 10 breeding sites. Collected mosquito larvae were reared under insectary conditions (22) and emerged adults (3 to 5 days old) were used for sugar choice experiments. All larvae collected in the wild turned out to be *An. gambiae* s.l. after rearing in the insectary. Thus, only *An. gambiae* s.l. were used for the sugar source comparison study.

Different fruits, identified as potential sugar sources based on previous studies elsewhere (20,23) and based on their prevalence in the study area and thus possible importance as a sugar source during the study period, were used in the sugar choice experiments. Fruits used in the experiments were guava (*Psidium guajava)*, melon (*Cucumis melo)*, banana (*Musa sp*), mango (*Mangifera indica*), marula *(Sclerocarya birrea)* and sugarcane (*Saccharum officinarum*).

A total of 810 insectary reared *Anopheles gambiae* s.l. were used for the sugar attractiveness experiments. Before exposure, mosquitoes were starved for 24 hours, given water only. Mosquito attractiveness to six naturally available fruits were determined compared to a control (distilled water) using an olfactory driven behavioural assay box. The assay was locally designed based on (21,24) with modifications to simultaneously compare the attractiveness of three different compounds in laboratory conditions.

Mosquito attractiveness experiments were conducted using nine experimental set-ups, each consisting of three experiments where fruit types were rotated every 30 minutes to minimize positional bias. Thirty fresh mosquitoes were released per experimental rotation, and a Latin square design was used to control for treatment effects and positional variations.

Each set-up included two fruit types and a control (e.g., mango, guava, and a control in tubes A, B, and C, respectively) (Figure. 2). Mosquitoes were introduced into a box olfactometer via a release chamber equipped with a mesh cover and sliding glass for controlled entry. Mini fans facilitated airflow, directing fruit odours toward the mosquitoes.

**Figure 2.**
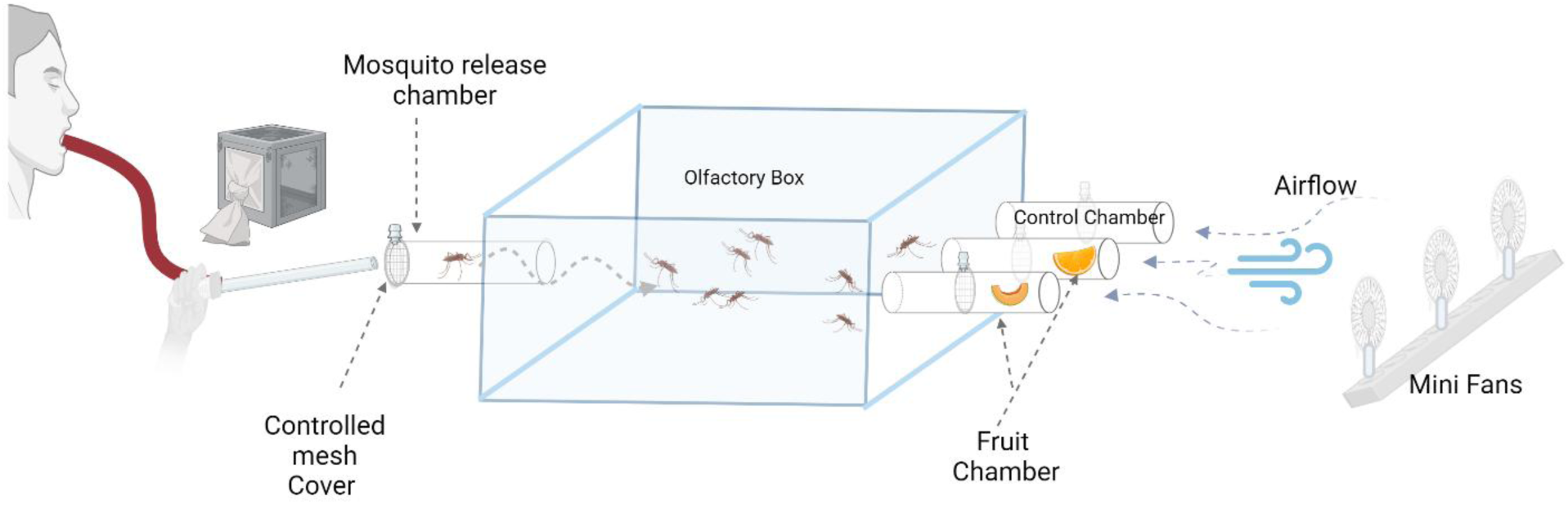
The olfactory driven behavioural assay box: An experimental set up which includes two types of fruits and a control. Starved *An. gambiae* s.l. from a cage were released through the release chamber into the olfactory box using a mouth aspirator. Release chamber was closed using a controlled mesh cover to hold mosquitoes. Mini fans were used to push scents into the box.

**Figure 3.**
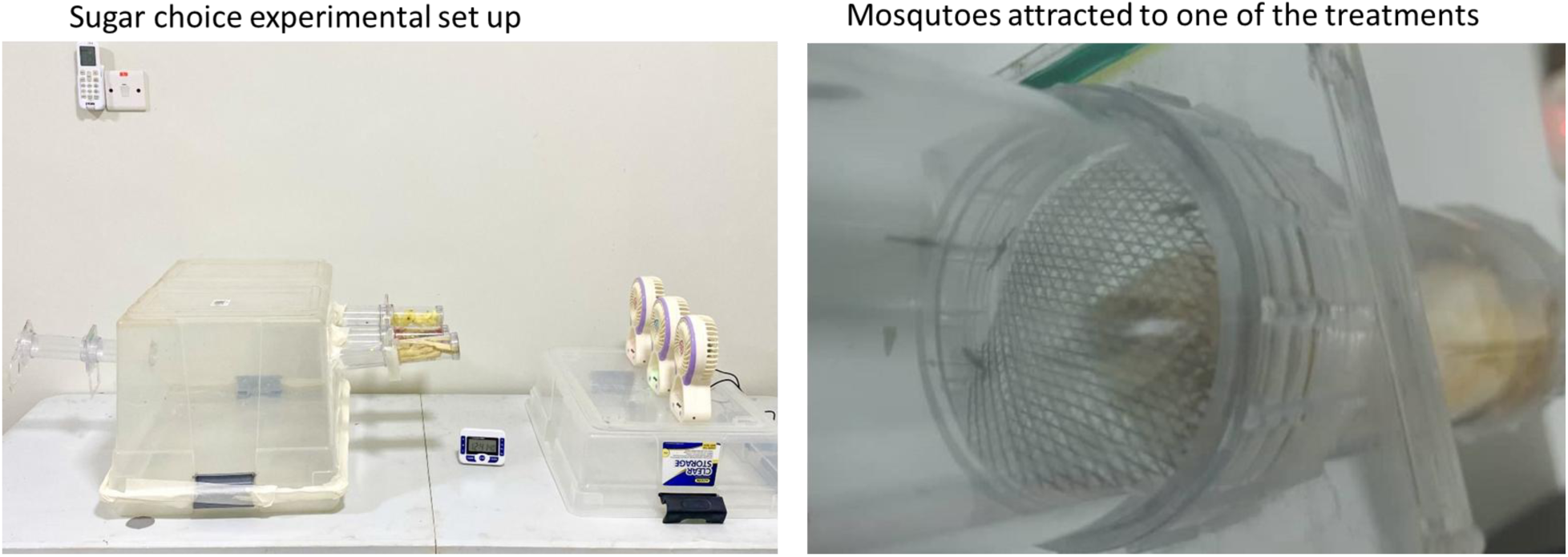
Sugar choice experiments. **A**. Mosquito sugar choice experimental set up. **B**. Mosquitoes attracted in a tube with one of the fruit treatments. Number of mosquitoes attracted by each treatment was determined by counting how many mosquitoes entered each tube with the different treatments compared to the control.

To reduce positional bias, treatment positions were switched twice across three experimental runs: first, with mango, guava, and control in tubes A, B, and C; then, with guava, control, and mango; and finally, with control, mango, and guava. The experiment was repeated for all set-ups. Between trials, acetone was used to clean the tubes to eliminate residual odours. Mosquito responses were recorded based on the number entering each treatment tube compared to the control.

### Statistical analysis

To assess the presence of sugars, we fitted a logistic regression model to estimate the presence of fructose in the wild caught anopheline mosquitoes. Due to the nested nature of the data, we included the trap as the random effect to investigate any possible differences between the traps.

To determine the preference for potential sugar sources for *An. gambiae s.l.,* we first performed an unadjusted analysis to compare the performance of the different treatments (sugar sources) in attracting mosquitoes. To achieve this, we performed the Kruskal-Wallis test for comparing the median number of mosquitoes across all the treatments. We then performed multiple comparison using Bonferroni correction to compare the number of mosquitoes caught by different pairs of treatments. Then in the adjusted analysis, we fitted a negative binomial regression model with the number of mosquitoes caught by each method as the response variable and treatment as a fixed effect. The negative binomial model was chosen to control the possible effect of overdispersion which occurs when the variance is greater than the mean. The treatment fixed effect allowed us to compare the different treatments. We fitted two models. In the first model, we did not account for the study design, thus leading to a simple negative binomial generalized linear model (GLM) without random effects. In the second model, we fitted a negative binomial mixed-effects model to account for the study design by letting the experiments be the random intercept. Model comparison was done through the Akaike Information Criteria (AIC) to select the best fitting for further inference. We then performed model diagnostics on the selected model.

Let 𝑌_𝑖_be the number of mosquitoes caught in treatment 𝑖. We assume that 𝑌_𝑖_ follows a Poisson distribution with mean 𝜇_𝑖_. The regression model with random effect is then expressed as

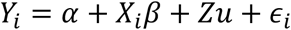

where 𝑿 is a design matrix of predictor variables, 𝜷 is a vector of fixed-effect coefficients and 𝒁 is the design matrix of random effects with a corresponding vector of random coefficients 𝒖. Lastly, 𝜖_𝑖_ is the residual error term. The fixed effect GLM without random effects does not have the term 𝑍𝑢. All analyses were done in the R environment for statistical computing.

## Results

### Exploring the presence of sugars in wild caught *Anopheles* mosquitoes

We evaluated the proportion of anopheline mosquitoes that fed on sugars using the cold anthrone test. A total of 181 out of the 187 *Anopheles* mosquitoes were successfully tested of which only 2 were males. Testing was conducted within two hours of collection on each collection day. Overall, over 40% (n=74) of the anopheline mosquitoes collected were positive for sugar, indicating they had fed on sugar sources. Logistic regression model further confirmed the presence of fructose in the wild caught anopheline mosquitoes (Table 1). There was no statistically significant difference on the presence of sugar among the four different *Anopheles* species. *An. gambiae* s.l. for example, exhibited an odds ratio (OR) of 1.75 (95% CI: 0.33-12.94) for sugar ingestion compared to *An. tenebrosus*, indicating a potentially, but statistically non-significant, higher likelihood of sugar consumption. In contrast, *An. funestus* (OR = 0.95, 95% CI: 0.16-7.70) and *An. coustani* (OR = 0.74, 95% CI: 0.11-6.17) showed lower odds of sugar ingestion relative to *An. tenebrosus*.

**Table 1:** Logistic regression model estimating the presence of fructose in the wild caught anopheline mosquitoes.

| Predictors | Number tested | Number positive | OR | CI | P |
| --- | --- | --- | --- | --- | --- |
| Intercept |  |  | 0.50 | 0.07 – 2.56 | 0.42 |
| *An. tenebrosus* | 6 | 2 | *Comparator* |  |  |
| *An. coustani* | 26 | 7 | 0.74 | 0.11 – 6.17 | 0.75 |
| *An. funestus* | 31 | 10 | 0.95 | 0.16 – 7.70 | 0.96 |
| *An. gambiae s.l.* | 118 | 55 | 1.75 | 0.33 – 12.94 | 0.53 |

Additionally, we determined the presence of sugar in mosquitoes against the type of trap used to collect them. Here, three traps; CDC light trap, Prokopack aspirator and barrier screen, which were used for mosquito collections were compared. From our findings, the CDC light trap and the barrier screen collected similar proportions of mosquitoes that fed on sugar (Figure 4). The results showed that 41.1 % of *Anopheles* mosquitoes collected by the CDC LT tested positive for sugar while 38.5 % collected by the barrier screen tested positive. For the barrier screens, it was interesting to note that mosquito abundance varied depending on the side of the barrier screen. Generally, over 93% (n=61) of the mosquitoes were collected from the side of the barrier screen facing the breeding sites while only a few (n=4) mosquitoes were collected from the other side of the barrier screen that faced people’s dwelling places. The Prokopack aspirator collected very few mosquitoes in general compared to the other collection methods (n=4). However, proportionally, the Prokopack had the highest percentage of mosquitoes that fed on plant sugars (75%-thus 3 out of the 4 mosquitoes had sugar in the gut) (Figure 4).

**Figure 4.**
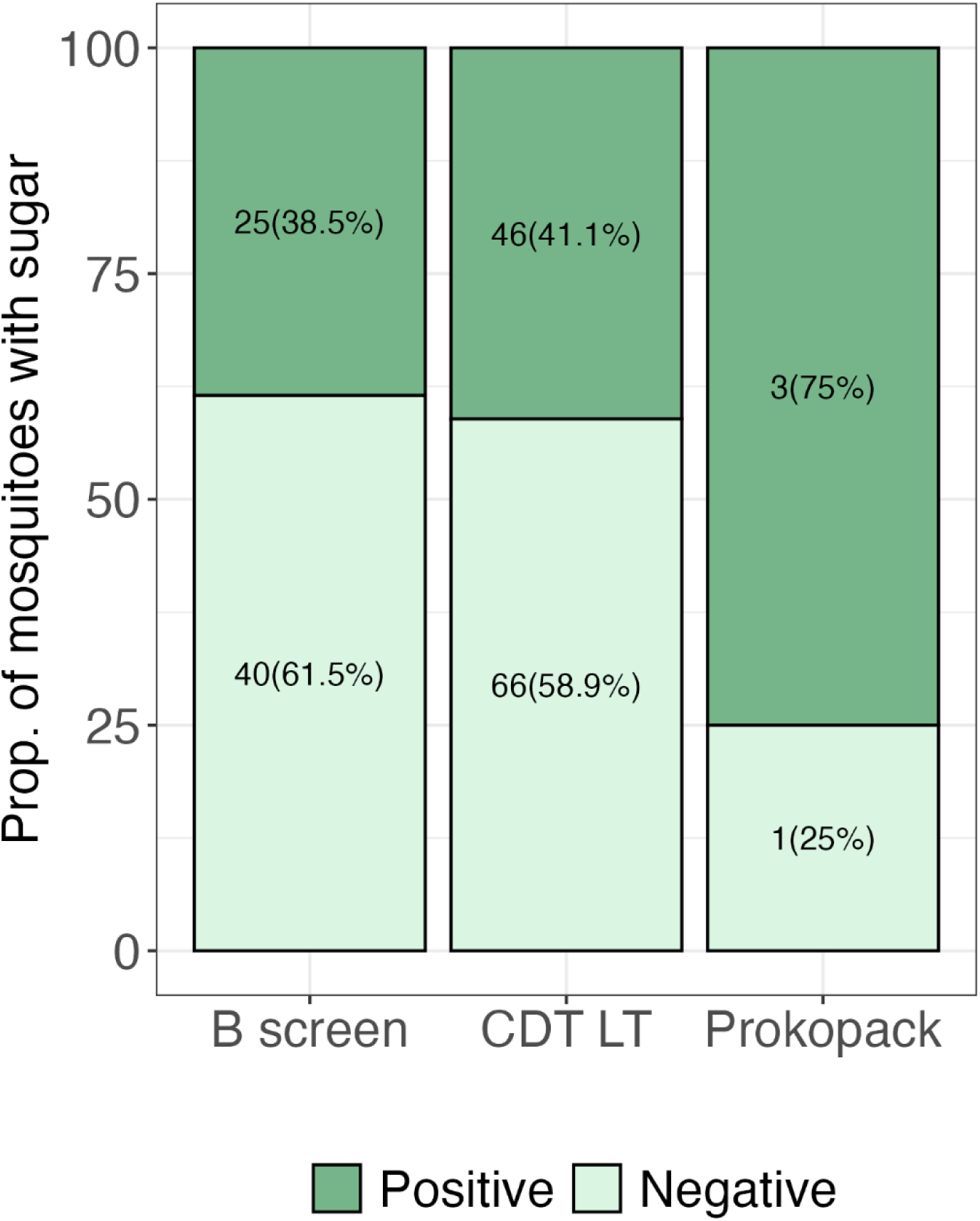
Proportion of mosquitoes that fed on plant sugars by trap type used. 75% of Anopheles mosquitoes collected by the Prokopack aspirator fed on sugar, 41.1 % of Anopheles mosquitoes collected by CDC LT fed on sugar while 38.5 % of Anopheles mosquitoes collected by barrier screen fed on sugar.

### Determining the attractiveness of fruits to *Anopheles gambiae* s.l. mosquitoes

We determined the attractiveness of six locally available fruits to anopheline mosquitoes in a laboratory setting. Figure 5 summarizes the average number of *Anopheles gambiae* s.l. attracted to local fruits. A total of 9 experimental set ups exposing 30 mosquitoes to each fruit were run. In general, the results show that guava caught the highest number of mosquitoes with marula being the least attractive fruit.

**Figure 5:**
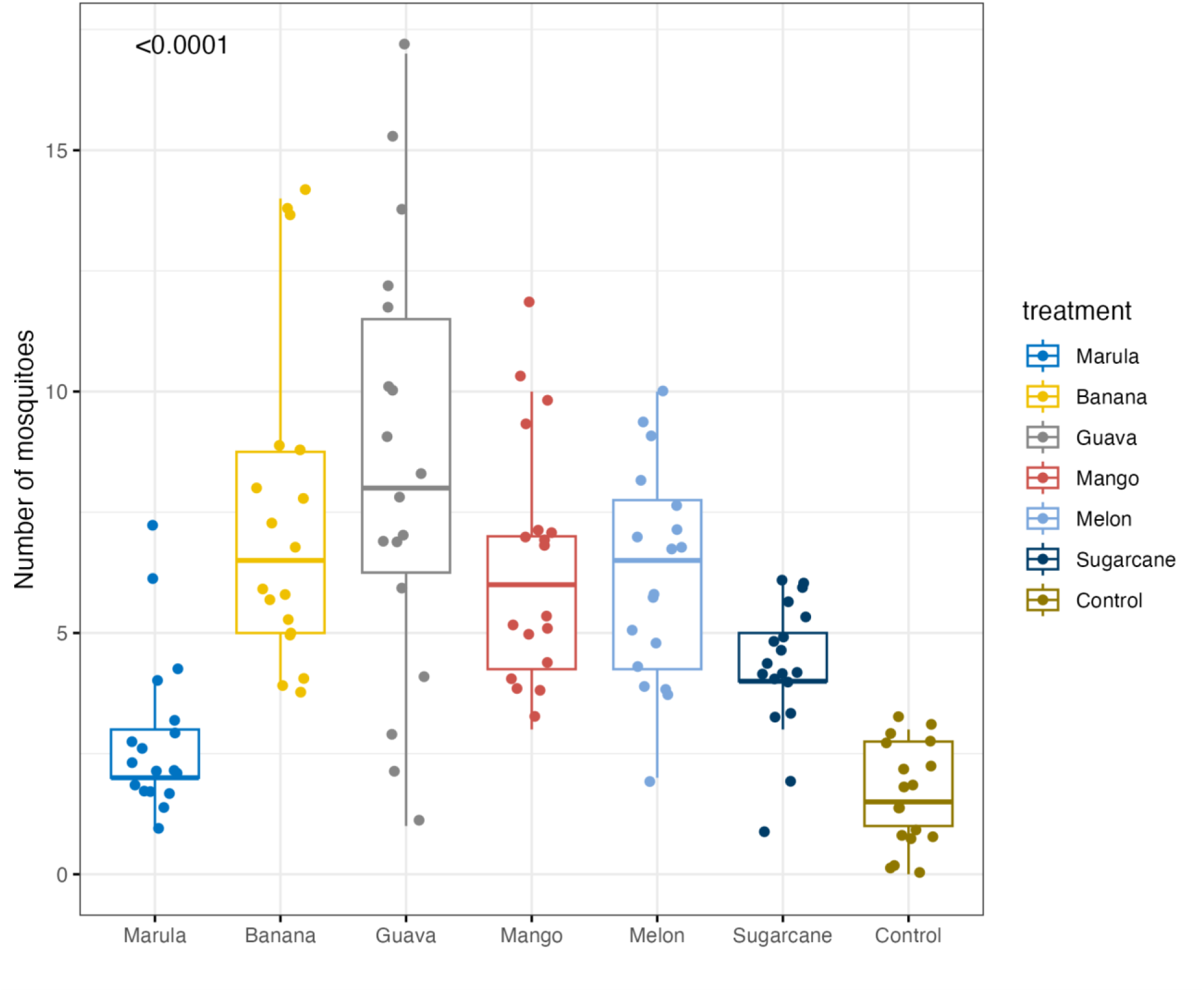
Attractiveness of mosquitoes to treatments in all experiments (experiments 1, 2 and 3) across the 11 set ups. Overall, the figures show that guava had the highest attractiveness with the highest median number of mosquitoes.

A Kruskal-Wallis test revealed significant differences in mosquito attractiveness among the treatments (p < 0.01). Pairwise comparisons demonstrated that marula differed significantly from banana, guava, mango, and melon (p < 0.001), but not from sugarcane. Banana did not significantly differ from guava, mango, or melon, but was significantly different from sugarcane (p = 0.02). We further fitted a GLM to compare fruit attractiveness to mosquitoes taking into account the nested random effects. Our analysis revealed a significantly higher incidence of mosquito attraction for all tested fruits compared to sugarcane (as reference). Table 2 shows the model results. Guava was the most attractive fruit as it attracted mosquitoes at approximately twice the rate of sugarcane (IRR = 1.97, 95% CI: 1.49-2.62). Banana attracted mosquitoes at a rate 1.68 times higher than sugarcane (IRR= 1.68, 95% CI: 1.26-2.24). Mango, and melon also demonstrated significantly higher mosquito attraction rates compared to sugarcane (IRR= 1.49, 95% CI: 1.11-2.01) and (IRR=1.45, 95% CI: 1.08-1.96) respectively. As expected, the control demonstrated significantly lower compared to the reference (IRR=0.14, 95% CI: 0.09-0.22) (Table 2). Comparing the attractiveness across treatments, mosquito attractiveness varied significantly between treatments (p<0.0001). Overall, guava was shown to be the most attractive treatment with the highest median mosquito count while marula and sugarcane were the least attractive (Figure 5).

**Table 2:** Comparison of fruit attractiveness to mosquitoes.

| Predictors | IRR | CI | p |
| --- | --- | --- | --- |
| Intercept | 1.07 | 0.85 – 1.35 | 0.568 |
| Sugarcane | *Comparator* |  |  |
| Marula | 1.32 | 0.92 – 1.90 | 0.130 |
| Banana | 1.68 | 1.26 – 2.24 | **<0.001** |
| Guava | 1.97 | 1.49 – 2.62 | **<0.001** |
| Mango | 1.49 | 1.11 – 2.01 | **0.008** |
| Melon | 1.45 | 1.08 – 1.96 | **0.014** |
| Control | 0.14 | 0.09 – 0.22 | **<0.001** |

## Discussion

We assessed the prevalence of sugar feeding in wild anopheline mosquitoes and the attractiveness of local fruits to *An. gambiae* s.l. in a laboratory setting. For the first time in Malawi, our study revealed what could be some of the potential plant sugar sources that are attractive to *Anopheles* mosquitoes.

Using the cold anthrone test, our study confirmed the presence of sugar in the guts of both primary and secondary malaria vectors in southern Malawi. Interestingly, the positive control (table sugar) initially turned blue green but reverted to yellow within minutes after testing, while fruit samples retained the blue-green colour for over 40 minutes. This may be due to the cold anthrone test being specifically designed to detect fructose, which is low in table sugar (25). Further studies are needed to confirm this in Malawi.

Different trapping methods were used to maximize collections and to capture mosquitoes at various physiological stages (e.g., host-seeking, resting, or dispersing). Our study found no significant differences in the proportion of sugar-fed mosquitoes across collection methods as both barrier screens and CDC light traps had similar proportions of sugar-fed mosquitoes. Although the Prokopack collected the highest proportion of sugar-fed mosquitoes, the lower number of mosquitoes collected by this method (n=2) increases the likelihood that the observed differences were due to chance.

The differences in trap performance were likely due to several factors. The CDC light traps for example, primarily attract host-seeking females, while barrier screens are more effective at intercepting host-seeking or dispersing female mosquitoes. On the other hand, the Prokopack is designed to collect resting mosquitoes of both sexes (26,27). This could in part explain why all males were collected by the Prokopack, although only a few males were collected.

The limited number of males collected during the study could be due to the timing of collections (6 PM–6 AM), the naturally low male abundance, or the inefficacy of trapping methods for males. Since males rely exclusively on sugars (28), their likelihood of being sugar-fed is high. However, the small sample size in this study prevents definitive conclusions about sex-specific sugar-feeding behavior.

Compared to a study in Kenya, where about 15.7 % of collected mosquitoes had fed on sugar (20), our study found that over 40% of mosquitoes had sugar in their guts. This difference may possibly be due to variations in mosquito species, local vegetation, or environmental conditions, highlighting the difference in suitability of tools that target sugar feeding and the need for more research on sugar-feeding behaviour across different ecological settings.

Having determined the mosquito sugar feeding behaviour, we further explored the attractiveness of different fruits to *An. gambiae* s.l. Using the box olfactometer behavioural bioassay, we found there was a significant attraction of *An. gambiae* s.l. to all fruit types tested compared to the control. Although all six fruits tested were attractive, there were variations on their attractiveness to *Anopheles gambiae* s.l., signifying that different plants may have differences in their importance as a sugar source. Elsewhere, it was also shown that *An. gambiae* s.l. demonstrates selectivity on its foraging activity towards plants species (29). In our study, guava, followed by banana seemed to be the most attractive while marula and sugarcane were the least attractive. However, bananas and guavas exhibited a greater variability in attractiveness, as indicated by their wider interquartile ranges (Figure 5), suggesting the presence of potential outliers in the data. Consistent with a previous study in Mali, guava was reported as the most attractive fruit to *An. gambiae* s.l. compared to other fruits (banana, sugarcane, melon) (30). In contrast to our findings the authors found bananas to be less attractive to *An. gambiae* s.l. while sugarcane was reported as one of the most attractive plants in the same study in Mali (30). The differences between our observations and the study in Mali could be due to differences in mosquito species. The predominant species identified in the current study was *An. gambiae* s.l. Although mosquito molecular identification was not performed in this study, recent studies suggest that *An. arabiensis* is likely the dominant species, where it accounts for over 80% of *An. gambiae* s.l. population in Chikwawa district (17,31–33). However, *An. gambiae* s.s. makes up about 86% of the *An. gambiae* s.l. found in the study area in Mali (23).

It Is also possible that the sugarcane types used in our experiment and the Mali study were different variants. Some studies have shown that the genotype and environment affect productivity and overall sugar content of sugarcane (34). Furthermore, sugar content of sugarcane can be influenced by the ages - whether they are early or mid-late maturing sugarcane varieties (35). These differences could further lead to differences in attractiveness to different in mosquitoes.

Sugar source preference can vary between species and seasons. Previous studies have shown that female mosquitoes exhibit preferences for specific plant sugars, with some plants being over 130 times more attractive than others (36–38). However, in the absence of preferred plants, mosquitoes can switch to feeding on less attractive but available alternatives (39). Therefore, plants that are abundant or whose flowering period coincide with peak density of mosquito populations could be explored for further evaluations.

In the current study, sugarcane was among the least attractive sugar sources for mosquitoes. However, the study area has one of the major sugar producing estate, home to the largest sugarcane plantations in southern Africa (17). Despite its lower attractiveness in this setting, the widespread presence of sugarcane could still influence malaria control efforts, as it may serve as an alternative sugar source when preferred and most attractive plants are scarce. This underscores the need for further research to better understand how sugar source availability influences vector populations across different ecological settings.

In general, the study showed that sugar feeding plays an important role in the survival of female malaria vectors, maintaining the survival rates of mosquitoes in the wild. There is clear preference for certain sugar sources by the *An. gambiae* s.l. tested, highlighting an opportunity to tailor tools to local preferences and lure them away from other sugar sources. The current study thus, adds evidence to studies from elsewhere (11,20,23,40,41) that natural sugar foraging behaviour in mosquitoes could be an important driver of mosquito abundance. This is because sugar is a good source of energy for mosquitoes and other insects and helps to sustain mosquito survival. Plant feeding has been attributed to *An. gambiae* survival by previous studies (42,43). The survival rate of mosquitoes is a very important factor on the mosquito vectorial capacity (42,44,45) and thus, sugar feeding plays a crucial role in mosquito life and could influence mosquito control. This is an important reason why novel mosquito control tools that specifically exploit the sugar feeding behaviour are important. Thus, studies on the effects of interventions on the survival rates of mosquitoes (i.e. sugar based control tools) are important and have previously been recommended.

This data could be significant in designing and deploying future studies and guide when selecting potential plant sugars to use in sugar baits when implementing sugar-based mosquito control tools in Malawi. However, since our sugar attractiveness study was laboratory based, there is need for field-based studies to replicate these findings in the real natural world. Another factor worth investigating is the role of visual stimuli in mosquito attraction to sugar sources. Previous research (29) has highlighted the importance of visual cues, showing that mosquitoes may be drawn to a plant based on its presentation. Another study, aimed at assessing sugar feeding behaviour of *Anopheles* mosquitoes in a setting with diverse natural sugar sources is ongoing in the study area and will be reported separately.

Additionally, the observation from the cold anthrone test, that a considerable proportion of the mosquitoes entering the community had already fed on natural sugar, as inferred from the barrier screen collections, has implications for the deployment of sugar baited trap stations. We hypothesize that deploying multiple bait stations around breeding sites or at the point of entry into communities might improve its efficiency than currently reported. While the practicality of this maybe questioned it may be worth exploring in separate studies.

## Limitations

One of the limitations of this study was the low number of mosquitoes collected, which may have been influenced by seasonality and the timing of the study. The study was conducted at the beginning of the dry season and did not align with peak mosquito abundance. Further studies, including both seasons, could increase opportunities to capture more mosquitoes and have comparisons across different seasons.

Additionally, the study was not able to analyze plant DNA from the wild caught mosquitoes. Thus, genetic sequencing of collected mosquitoes could complement the cold anthrone tests and sugar choice experiments. This could have confirmed whether mosquitoes had fed on the plants identified as attractive in behavioural bioassays. Future studies should incorporate molecular analysis of mosquito gut contents to verify plant feeding behaviour in field conditions.

While the composition of plant sugar may influence mosquito attraction, our study did not analyse the sugar content of the plants used. Further research should investigate chemical composition of these sugars for better understanding of their role in mosquito attraction. A recent study (46) has demonstrated the potential to identify plant hosts from nectar metabolites inside mosquito guts, and a similar study would be valuable in Malawi.

## Conclusion

Sugar feeding is a key activity for *Anopheles* mosquitoes and presents a potential target for control efforts. In this setting with abundant sugarcane, guava was identified as the most attractive sugar source for *An. gambiae* s.l., followed by banana, mango, and melon, with sugarcane being the least attractive. Understanding local sugar source preferences can help tailor novel intervention strategies to specific environmental contexts.

## DECLARATIONS

### Ethics approval and consent to participate

The study was conducted in Malawi, with an ethical approval from the College of Medicine Research and Ethics Committee (COMREC) (P.05/23/4101). Prior to the study commencement, support was obtained support from the District Health Officer of Chikwawa and officials from the Illovo sugar estate. No human participants were recruited in this study and thus, no informed consenting was required.

### Consent for publication

Not applicable

### Availability of data and materials

The datasets used and/or analysed during the current study are available from the corresponding author on reasonable request.

### Authors’ contributions

KZ: conception, data collection, data analysis, and writing the first draft of the manuscript, SK: data collection, writing and editing; JC: data analysis, writing and editing; RM: data collection, writing and editing; JT: supervision, writing and editing. All authors contributed to the writing of the manuscript and reviewed the final draft.

## Acknowledgements

We are very grateful to Dr. Christopher M. Jones from Liverpool School of Tropical Medicine for his support during the study funding application. Special thanks to the Illovo sugar estate community for accepting us to conduct this study in their area.

